# An early decline in ETPs reflects fewer pre-thymic progenitors and altered signals from the thymus microenvironment

**DOI:** 10.1101/2022.01.18.476832

**Authors:** Jayashree Srinivasan, Anusha Vasudev, Hilary J. Selden, Encarnacion Perez, Bonnie LaFleur, Shripad A. Sinari, Andreas Krueger, Ellen R. Richie, Lauren I. R. Ehrlich

**Affiliations:** Department of Molecular Biosciences, The University of Texas at Austin, Austin, TX; Department of Epigenetics and Molecular Carcinogenesis, The University of Texas MD Anderson Cancer Center, Houston, TX; Center for Biomedical Informatics and Statistics, The University of Arizona, Tucson, AZ; Institute for Molecular Medicine, Goethe University Frankfurt am Main, 60590 Frankfurt am Main, Germany; Department of Oncology, Livestrong Cancer Institutes, The University of Texas at Austin Dell Medical School, Austin, TX

**Keywords:** Thymus involution, early T-cell progenitors, hematopoiesis, aging

## Abstract

Age-related thymus involution results in decreased T-cell production, contributing to increased susceptibility to pathogens. Elucidating mechanisms underlying involution will inform strategies to restore thymopoiesis. The thymus is colonized by circulating bone marrow (BM)-derived thymus seeding progenitors (TSPs) that differentiate into early T-cell progenitors (ETPs). We find ETP cellularity declines as early as 3 months (3MO) in mice. This initial ETP reduction could reflect changes in pre-thymic progenitors and/or thymic stromal niches. We demonstrate that the number of functional TSP/ETP niches is not reduced with age. Instead, the number of pre-thymic BM and circulating lymphoid progenitors is substantially reduced by 3MO, although their intrinsic ability to seed and differentiate in the thymus is maintained. Additionally, Notch signaling in BM progenitors and ETPs declines by 3MO, indicating defective niche quality contributes to the reduction in ETPs. Together, these findings indicate that diminished BM lymphopoiesis and thymic stromal support contribute to the initial decline in ETPs, setting the stage for progressive thymus involution.

**Summary statement:** The number of early T-cell progenitors declines by 3 months of age in mice. This decline reflects a sharp drop in circulating thymus seeding progenitors, fewer bone marrow lymphoid progenitors, and reduced Notch signaling in both bone marrow and thymus.

## Introduction

T cells develop in the thymus which provides a unique and complex stromal microenvironment essential for T-cell lineage commitment, differentiation and selection (Han & Zuniga-Pflucker, 2021; Yui & Rothenberg, 2014). Since thymocytes do not have the ability to self-renew, the thymus periodically recruits bone marrow (BM)-derived thymus seeding progenitors (TSPs) which are essential to maintain T cell production throughout life (Krueger et al., 2017). TSP entry is a well-orchestrated multistep process mediated by the adhesion molecules P-selectin, ICAM-1, and VCAM-1, the chemokines CCL25 and CCL21, and other niche factors, like KITL, supplied by thymic endothelial cells (Buono et al., 2016; Krueger et al., 2010; Rossi et al., 2005; Scimone et al., 2006; Zlotoff et al., 2010). After entering the thymus via vasculature at the cortico-medullary junction, TSPs differentiate into highly proliferative CD4^-^ CD8^-^ CD44^hi^ c-kithi early T-cell progenitors (ETPs) in the thymic cortex. Commitment of developing thymocytes to the T-cell lineage depends on inductive signals provided by cortical thymic epithelial cells (cTECs) (Han & Zuniga-Pflucker, 2021; Petrie & Zuniga-Pflucker, 2007; Thompson & Zuniga-Pflucker, 2011; Yui & Rothenberg, 2014). In particular, activation of Notch signaling in ETPs via Delta-like Notch ligand 4 (DLL4) expressed by cortical thymic epithelial cells (cTECs) is indispensable for T-cell lineage commitment and differentiation (Hozumi, Mailhos, et al., 2008; Hozumi, Negishi, et al., 2008; Koch et al., 2008). Since ETPs give rise to all downstream T cell subsets, thymopoiesis and T cell development depend on both continued influx of TSPs into the thymus and the availability of functional stromal niches that support TSP entry and downstream ETP differentiation.

During age-associated thymus involution, thymus size and cellularity undergo a progressive decline accompanied by reduced output of naïve T cells. Thymus involution is initiated early in life, by ∼7 weeks of age in mice (Chen et al., 2009; Hale et al., 2006). Thus, we first asked if ETP cellularity is adversely affected during the earliest stages of thymus involution. Here, we find that the decline in ETP cellularity is initiated early, between 1MO and 3MO of age in mice, and further declines with advancing age. Thymus involution involves not only a reduction in thymocyte cellularity, but also a decline in the number, composition, proliferative capacity, and organization of the TEC compartment (Baran-Gale et al., 2020; Chinn et al., 2012; Gray et al., 2006; Griffith et al., 2012; Lepletier et al., 2019). TEC degeneration is a key driver of age-associated thymic involution, as sustaining or restoring TECs prevents or reverses involution, respectively (Bredenkamp et al., 2014; Chen et al., 2009; Cheng et al., 2010; Garfin et al., 2013; Klug et al., 2000). Given that thymic stromal niche factors play a key role in recruitment of TSPs, and thymic stromal signals are required for ETP survival, proliferation, and T-lineage commitment, we anticipated the early age-associated decline in ETPs would be driven by changes in the thymic microenvironment.

Here, we used several approaches to determine whether the early loss of ETPs is due to age-related changes in availability of functional TSP/ETP niches and/or a decline in pre-thymic lymphoid progenitors. First, we used a multicongenic progenitor transfer approach to quantify the number of functional TSP niches from the onset of involution through middle-age (Zietara et al., 2015). These studies revealed that the number of TSP niches is sustained, despite substantial involution, through 12MO of age. We thus considered whether the reduction in ETP cellularity could be due to a decline in the hematopoietic progenitors that seed the thymus. Surprisingly, we observe a significant decline in the number of lymphoid progenitors with T-lineage potential in the BM and blood at 3MO of age, although their cell-intrinsic ability to seed and differentiate in the thymus remains intact. Because Notch signaling plays a key role in promoting T-lineage specification and progenitor differentiation, we also interrogated niche function by evaluating *Dll4* expression by cTECs and assessing Notch signaling activity in ETPs and BM progenitors. Notably, we find reduced Notch signaling in T-lineage precursors in both the BM and thymus by 3MO of age. Collectively, these findings demonstrate a surprisingly early decline in BM lymphoid progenitors and circulating TSPs, along with diminished Notch signaling in the BM and thymus microenvironments, implicating both pre-thymic and thymic changes in the early decline of ETP cellularity at the outset of thymus involution.

## Results and discussion

### ETP and total thymocyte cellularity decline by 2MO of age

We first quantified age-associated changes in ETPs and downstream thymocyte subsets between 1 and 12MO of age (Fig. 1 and Fig. S1 A). Total thymus cellularity declines with age, dropping significantly by 2MO, consistent with previous reports that thymus involution begins early in life (Fig. 1 A) (Chen et al., 2009; Hale et al., 2006; Lepletier et al., 2019). Notably, ETP cellularity also declines progressively, starting at 1.5MO, which precedes, but otherwise mirrors the reduction in total thymocytes (Fig. 1 A). There is a strong correlation between ETP and total thymocyte numbers through 12MO of age (Fig. 1 B), and this correlation extends to subsequent thymocyte subsets (Fig. S1 B), showing that increased numbers of downstream thymocytes do not compensate for the loss of ETPs with age. Moreover, the consistent DN2 to ETP ratio over the first year of life further suggests that ETP differentiation is not impaired with age (Fig. 1 C).

**Figure 1.**
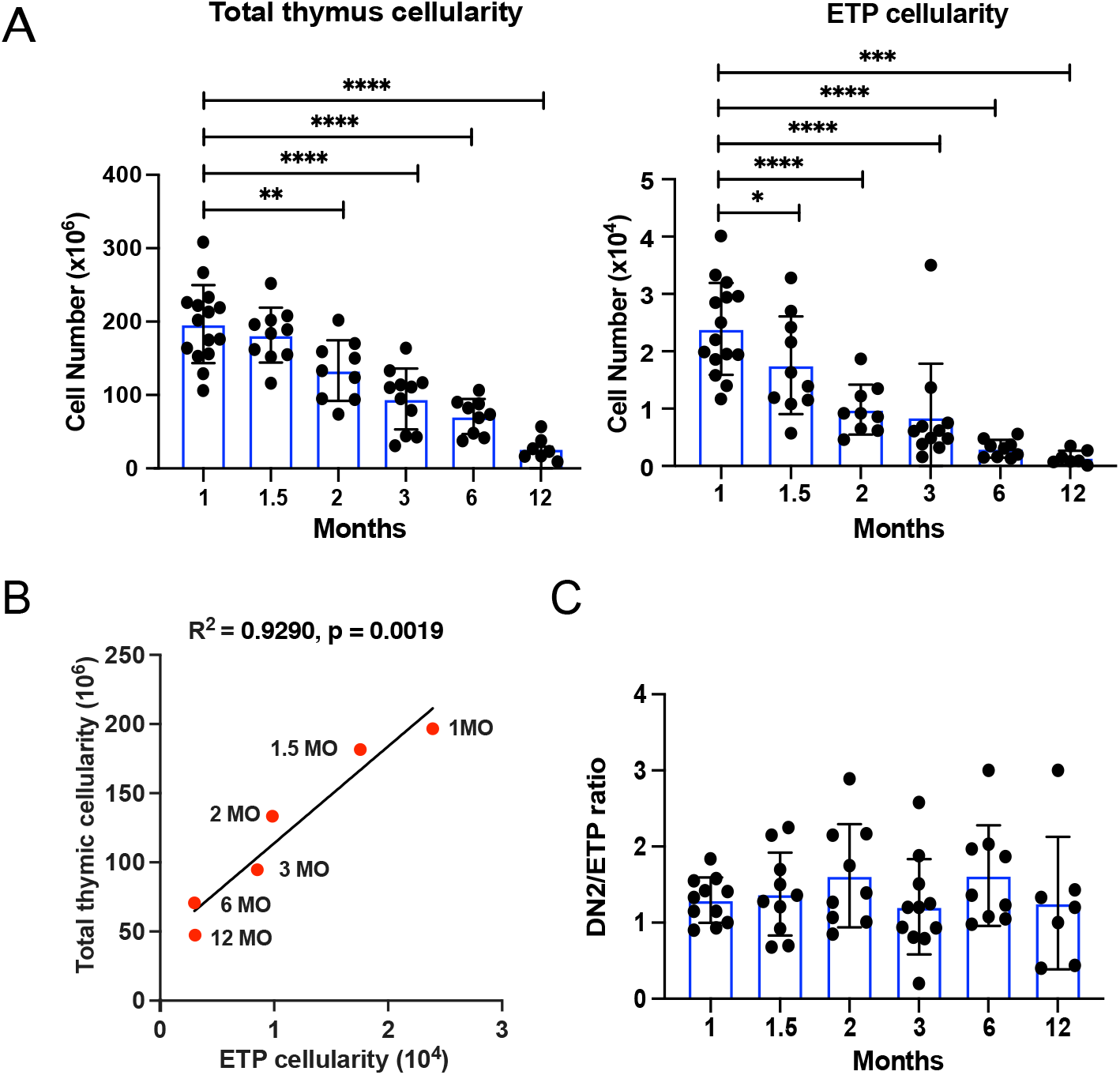
ETP and total thymocyte cellularity declines by 2MO of age. (A) Quantification of total thymocytes and ETPs in C57BL/6J mice at the indicated ages. (B) Linear regression analysis of total thymic cellularity and ETP cellularity at each age. R^2^ = coefficient of correlation (C) Ratio of DN2 to ETP cellularity at each age. Symbols represent A) and C) data from individual mice and B) an average of 7-14 mice at each age. Bars represent means +/-SEM from 3-4 independent experiments for each age group. Statistical analysis in (A) was performed using one-way ANOVA with Dunnett’s multiple comparisons test where *p<0.05, **p<0.01, ***p<0.001, ****p<0.0001. See also Fig. S1. Gating strategy for identification of thymocyte subsets and correlation of ETP cellularity with downstream thymocyte subsets.

### The number of available thymic niches for TSPs does not decrease with age

The age-associated decline in ETP cellularity could be a function of changes in the thymic stromal microenvironment and/or defects in pre-thymic lymphoid progenitors. Given the important role of the thymic microenvironment in supporting all stages of developing T cells and its known impact on involution, we first tested whether the reduction in ETPs is due to an age-related decline in the number of functional progenitor niches that support the early stages of thymopoiesis. We used a modification of a previously reported multicongenic progenitor transfer approach to quantify available, functional TSP niches from 1MO-12MO of age. This approach previously revealed that a young thymus contains ∼10 available niches that can be colonized by thymocyte progenitors at any one time (Zietara et al., 2015). We FACS sorted Lin^-^ Flk2^+^CD27^+^ BM progenitors, which encompass all thymus seeding and T cell progenitor activity in the BM (Serwold et al., 2009), from 8 different congenic strains, mixed them at equal ratios, and transplanted them into non-irradiated recipient mice of different ages (Fig. 2 A and Fig. S2, A and B). Recipient thymuses were analyzed after 21 days to allow sufficient time for donor progenitors to complete one wave of T-cell differentiation (Serwold et al., 2009). To determine the number of missing congenic tags, we quantified the number of different donor strains present in each recipient thymus by flow cytometry (Fig. S2 C), and then used mathematical modeling to estimate niche numbers. If the number of available niches that support TSP seeding declines with age, then fewer donor strains will be detected in thymuses of older recipients. However, we found that the number of available, functional TSP niches does not decline with age; to the contrary there is a trend toward an increased number in 6 and 12MO recipients (Fig. 2, B and C). Therefore, the age-associated decline in ETP cellularity is not due to a reduction in the number of functional niches available to thymus seeding progenitors.

**Figure 2.**
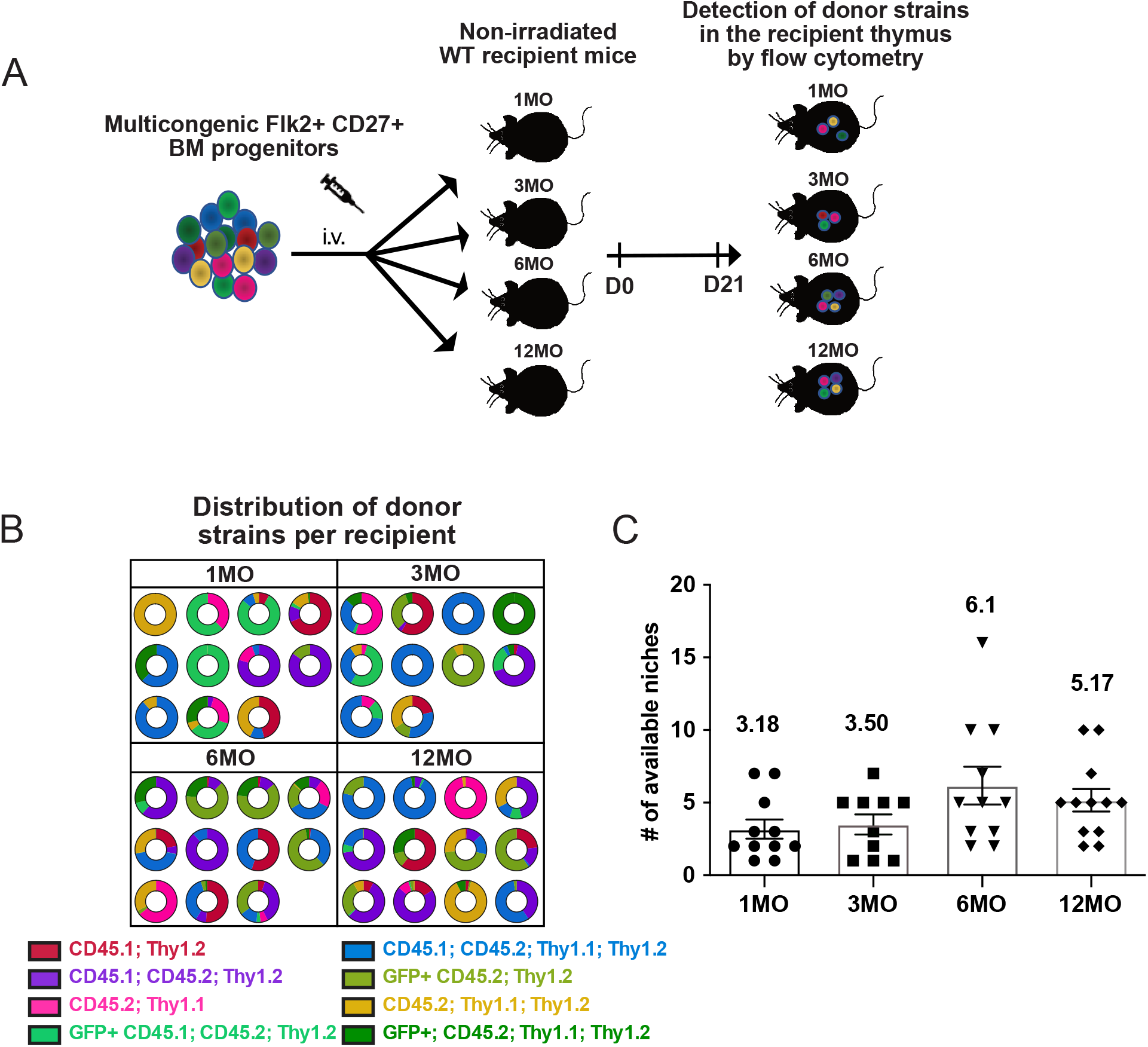
The number of available, functional TSP niches does not decline with age. (A) Schematic of the multicongenic barcoding experiment to quantify available niches in mice of different ages. Lin^-^Flk2^+^CD27^+^ BM progenitors were isolated from 8 different congenic strains (Fig. S2 B), mixed at equal ratios and i.v. injected into non-irradiated 1MO, 3MO, 6MO and 12MO mice. After 21 days, the thymi from recipient mice were analyzed by flow cytometry to quantify the number of detectable donor strains that underwent T-cell differentiation. (B) Pie charts illustrate the number and relative frequency of distinct color-coded donor strains in individual recipients of the indicated ages. (C) The estimated number of available functional niches in each age recipient as determined by multinomial sampling. Symbols indicate data from individual mice; bars represent means +/-SEM compiled from 4 independent experiments (n=10-12 mice per recipient age). See also Fig. S2 C. Identification of donor strain-derived thymocytes in recipient thymuses following transfer of multicongenic progenitors.

### 1MO and 3MO T-cell progenitors seed and thrive comparably in the thymus environment

We next asked whether the decline in ETPs reflects intrinsic defects in 3MO TSPs, compromising their ability to seed and differentiate in available thymic niches. Thus, we performed competitive heterochronic progenitor transfer assays in which isolated Lin^-^ Flk2^+^CD27^+^ BM progenitors obtained from 1MO and 3MO congenic donor strains were mixed at an equal ratio and injected i.v. into 1MO versus 3MO non-irradiated recipients. If BM progenitors from 3MO donors have intrinsic defects, then we would expect to detect fewer 3MO compared to 1MO donor-derived cells in recipient thymuses. However, comparable percentages and numbers of 1MO and 3MO donor cells were recovered from recipient thymuses of both ages (Fig. 3, A and B), indicating that TSPs from 3MO donors are fully competent to seed and thrive in the thymus. Moreover, there is no apparent defect in the differentiation capacity of 3MO TSPs, as a similar percentage of DP thymocytes are generated from 1MO versus 3MO donor cells (Fig. 3 C).

**Figure 3.**
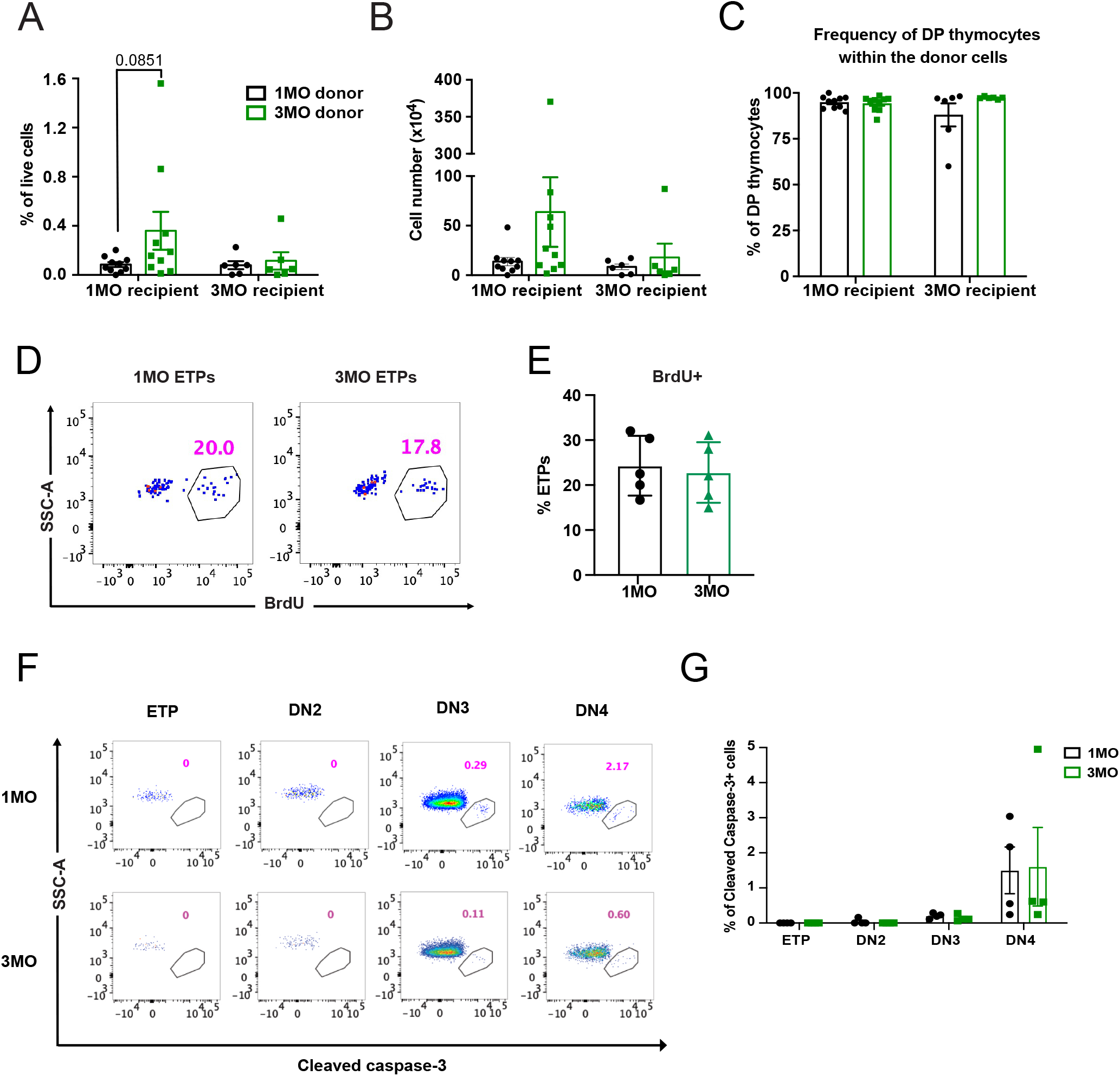
3MO T-cell progenitors show comparable seeding and differentiation capacity relative to 1MO progenitors. (A-C) Donor chimerism in 1MO and 3MO recipient thymuses 21d after transplantation of an equal number of 1MO and 3MO Lin^-^Flk2^+^CD27^+^ BM progenitors into 1MO or 3MO non-irradiated recipients. Quantification of the (A) frequency and (B) number of 1MO and 3MO donor-derived thymocytes in recipients of the indicated ages. (C) Quantification of the percentage of DP thymocytes derived from 1MO and 3MO donors detected in recipient mice of the indicated ages. Data are pooled from 4 independent experiments (n= 6-10 mice per recipient age). (D) Representative flow cytometry plots and (E) quantification of the percentage of ETPs that incorporated bromodeoxyuridine (BrdU) 8 hours after i.p. injection in mice of the indicated ages. Data are compiled from 4 independent experiments (n=5 mice per age group). **(F)** Representative flow cytometry plots and (G) quantification of the frequency of cleaved caspase-3^+^ cells in the indicated DN thymocyte subsets from 1MO and 3MO mice. Data are pooled from 4 independent experiments (n=4 mice per age group). (A-C, E and G) Symbols represent data from individual mice. Bars represent means +/-SEM. p value for (A) was obtained by two-way ANOVA with Sidak’s multiple comparison test. See also Fig. S3. Notch-Venus signaling reporter demonstrates faithful expression by immature thymocyte subsets.

We next considered the possibility that changes in proliferation or survival could account for the decline in ETPs at 3MO. However, a comparable percentage of 1MO and 3MO ETPs incorporated BrdU (Fig. 3, D and E), and analysis of intracellular cleaved caspase-3 showed that apoptosis is negligible in ETPs at both 1MO and 3MO of age (Fig. 3, F and G). Taken together, the data suggest that the decline in ETP numbers at 3MO of age is not a consequence of defects in the ability of 3MO thymus seeding progenitors to thrive in the thymus.

### Lymphoid progenitors with T-lineage potential decline in the BM and blood of 3MO mice

As we found no apparent defects in either the availability of functional TSP niches with age or the ability of 3MO TSPs to enter and thrive in those niches, we asked whether the number of BM or circulating lymphoid progenitors decreases by 3MO of age. A significant reduction in the number of TSPs could play a key role in the age-associated decline in ETP cellularity. To assess this possibility, we used flow cytometry to quantify circulating Lin^-^Flk2^+^CD27^+^ cells, which comprise all thymic seeding activity in the blood (Serwold et al., 2009) (Fig. 4 A). We also quantified upstream Lin^-^Flk2^+^CD27^+^ lymphoid progenitors in the BM (Fig 4 C). Interestingly, the frequency and number of Flk2^+^ CD27^+^ lymphoid progenitors in both blood and BM undergoes a significant decline between 1MO and 3MO of age (Fig 4, B and D). Flk2^+^ MPPs and Ly6d^-^ CLPs (Inlay et al., 2009) are the lymphoid progenitors with T-lineage potential within the Lin^-^ Flk2^+^CD27^+^ compartment (Serwold et al., 2009), and both of these subsets also decline significantly by 3MO of age (Fig. 4, E and F). These results demonstrate a surprisingly early reduction in the number of BM lymphoid progenitors and circulating TSPs, indicating that the decrease in ETP cellularity by 3MO is likely due, at least in part, to a reduction in the number of progenitors available to enter the thymus and differentiate therein.

**Figure 4.**
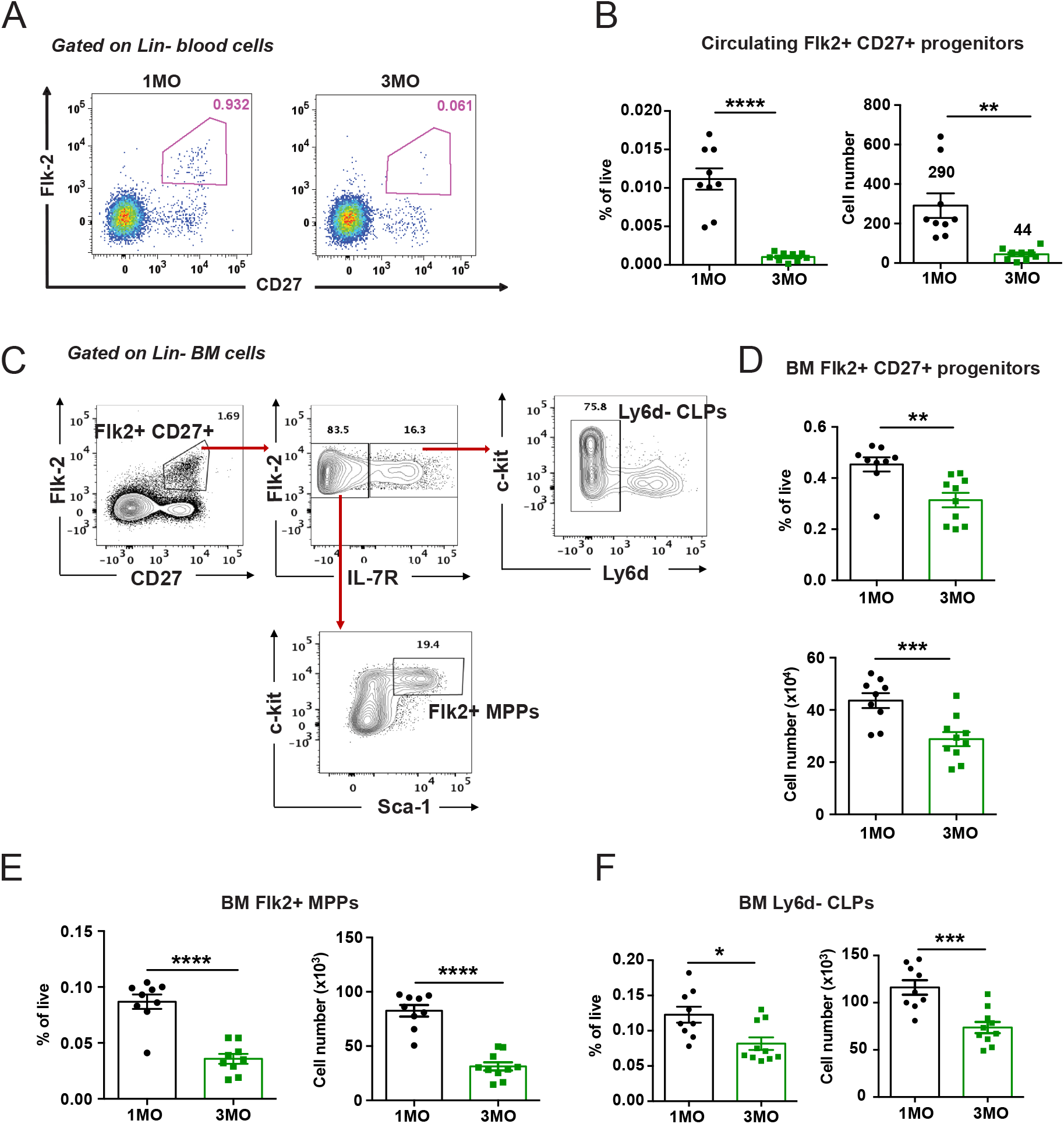
The number of circulating TSPs and BM lymphoid progenitors declines between 1MO and 3MO of age. (A) Representative flow cytometry plots showing the percentage of Flk2^+^CD27^+^ progenitors within the Lin^-^ compartment of the blood in 1MO and 3MO mice. (B) Frequency and absolute cell numbers of circulating lymphoid progenitors in the blood of 1MO and 3MO mice. (C) Representative gating strategy for quantification of Lin^-^Flk2^+^CD27^+^ progenitors, Ly6d^-^ common lymphoid progenitors (CLPs), and Flk2^+^ multipotent progenitors (MPPs) in the BM. (D-F) Frequency and absolute numbers of (D) Flk2^+^CD27^+^ progenitors, (E) Flk2^+^ MPPs and (F) Ly6d^-^ CLPs in BM of 1MO and 3MO mice. Symbols represent data from individual mice at each age. Bars represent means +/-SEM of data compiled from 3 independent experiments (n=9 mice per age group). Statistical analysis was performed using Student *t-*test *p < 0.05, **p <0.01, ***p<0.001, ****p<0.0001.

### Notch signaling is downregulated by 3MO in ETPs and BM lymphoid progenitors

Notch signaling in bone marrow lymphoid progenitors plays a critical role in differentiation of T-lineage progenitors (Chen et al., 2019; Tikhonova et al., 2019); thus, we considered the possibility that reduced Notch signaling could contribute to the decrease in BM lymphoid progenitors at 3MO of age. To test this hypothesis, we used the CBF-H2B-Venus strain in which expression of an H2B-Venus fusion protein is driven by Notch activation (Nowotschin et al., 2013). Due to perdurance of the H2B-Venus fluorescent signal, this reporter identifies cells that are currently undergoing or have recently undergone Notch signaling. Although the frequency of Notch-signaled Flk2^+^ MPPs and Ly6d^-^ CLPs was comparable at 1MO and 3MO (Fig. 5, A-D), there was a notable enrichment of Ly6d^-^ CLPs expressing low levels of the Venus reporter in 3MO relative to 1MO mice (Fig. 5, E and F). This suggests that by 3MO of age, Notch signaling wanes in Ly6d^-^ CLPs in the BM.

**Figure 5.**
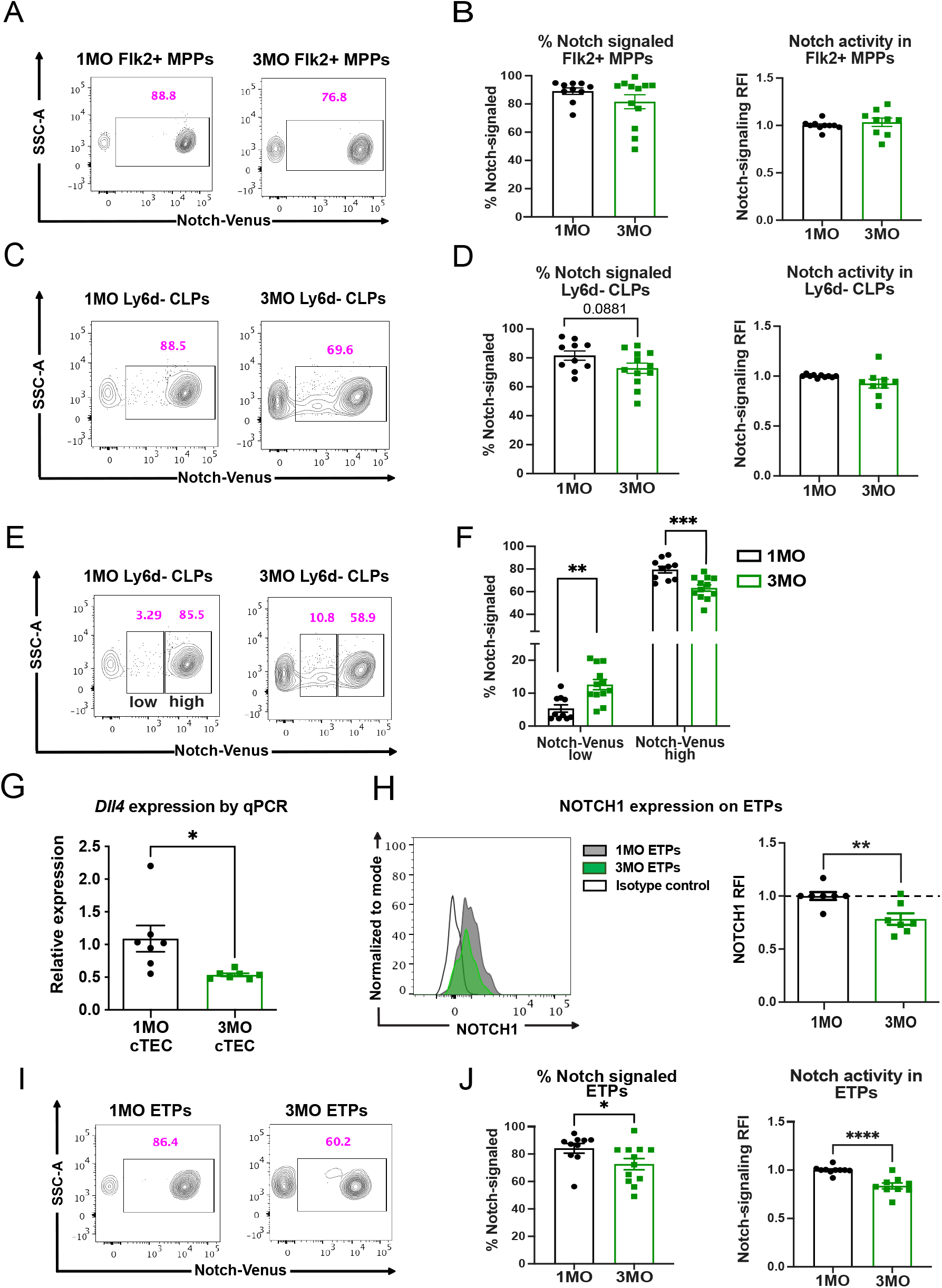
Age-associated changes in Notch signaling correlate with the decline in BM lymphoid progenitors and ETPs at 3MO. (A-D) Representative flow cytometry plots, as well as quantification of the frequencies and relative fluorescence intensities (RFIs) of Notch-Venus reporter expression in (A and B) BM Flk2^+^ MPPs and (C and D) BM Ly6d^-^ CLPs. (E) Representative flow cytometry plots showing gating to distinguish Notch-Venus low from Notch-Venus high cells. (F) Frequencies of Notch-Venus low and Notch-Venus high Ly6d^-^ CLPs in 1MO and 3MO BM. (A-F) Data are pooled from 5 independent experiments (n= 9-12 mice per age group). (G) qPCR analysis showing relative *Dll4* expression by EpCAM^+^ Ly51^+^ cTECs from 1MO and 3MO C57BL/6J mice. Data are pooled from 2 biological experiments with 3-4 technical replicates per experiment, and expression levels were normalized to those at 1MO of age. (H) Representative histogram (left) and quantification of RFIs (right) of NOTCH1 cell surface expression on thymic ETPs. Data are normalized to the average NOTCH1 MFI of 1MO mice in each individual experiment. Data are pooled from 3 independent experiments (n=7 mice per age group). (I) Representative flow cytometry plots and (J) frequencies and RFIs of Notch-Venus reporter expression in thymic ETPs. Data are pooled from 5 independent experiments (n= 9-12 mice per age group). (B, D, F, H and J) Symbols represent data from individual mice at each age and bars represent means +/-SEM. Statistical analysis was performed using Student *t-*test *p < 0.05, **p<0.01, ***p<0.001, ****p<0.0001.

cTECs provide a critical supportive niche for thymic ETP development, suggesting that age-associated deterioration of the TEC compartment could also contribute to the reduction in ETPs by 3MO of age. Activation of the Notch signaling pathway in ETPs is required for their development and T-cell lineage commitment. Our previously reported gene expression data show that expression of the Notch ligand *Dll4* is downregulated in cTECs between 1 and 3MO of age (Ki et al., 2014). Quantitative RT-PCR analysis of isolated 1MO and 3MO cTECs confirms reduced *Dll4* expression by 3MO of age (Fig. 5 G). In addition, the level of NOTCH1 expression on ETPs declines significantly by 3MO (Fig. 5 H). To determine if the decreased levels of NOTCH1 on ETPs and reduced *Dll4* expression by cTECs at 3MO of age are associated with diminished Notch signaling in ETPs, we first tested whether the Notch-Venus reporter is faithfully expressed in immature thymocyte subsets known to undergo Notch-signaling. As expected, reporter levels were high in ETPs and DN2s and dropped in the DN3 and DN4 subsets (Fig. S3). Notably, the frequency of ETPs undergoing Notch signaling declines between 1MO and 3MO of age (Fig. 5, I and J). Also, the lower relative mean fluorescence intensity (RFI) of the Venus reporter in 3MO ETPs indicates a reduced level of Notch signaling. Taken together, these data indicate that the early decline in ETP cellularity is associated with diminished Notch signaling activity in T-lineage progenitors in both the BM and thymus microenvironments.

It is well established that thymus cellularity and T-cell output decline during age-associated thymus involution (Chinn et al., 2012). While prior studies focused on comparative analysis of thymuses from young mice (1-4MO of age) versus severely involuted thymuses from middle-aged and old mice (9-28MO of age), the present investigation focuses on changes that occur during the initial stages of thymus involution between 1 and 3MO of age. In agreement with other reports (Chen et al., 2009; Hale et al., 2006; Lepletier et al., 2019), we find a significant decline in thymus cellularity by 3MO of age. Furthermore, the progressive reduction in total thymocyte numbers with age is highly correlated with a reduction in the number of ETPs, which contain progenitors that give rise to all subsequent stages of developing T cells. These results indicate that the early decline in ETPs plays a key role in initiating age-associated thymus atrophy. Given that TSPs, ETPs and all subsequent stages of developing T cells rely on inductive and supportive signals from the thymic microenvironment, which undergoes profound age-associated changes that drive thymus involution, we first tested whether a reduction in the thymic niches that support TSPs and ETPs are diminished by 3MO of age. Unexpectedly, we did not find an age-related decline in the number of functional, available TSP niches between 1 and 12MO of age. In fact, there was a slight increase in these niches by 12MO of age. Thus, we subsequently investigated whether the early decline in ETP numbers reflects a reduction in BM and/or circulating T-cell progenitors. Our studies reveal a dramatic decline in circulating Flk2^+^CD27^+^ progenitors, which contain TSPs, as well as a reduction in BM lymphoid progenitors by 3MO of age. Furthermore, expression of the Notch ligand DLL4 by cTECs is reduced by 3MO, as is Notch signaling activity in ETPs and BM-resident CLPs. Overall, our results suggest that both a decline in pre-thymic lymphoid progenitors in the BM and blood, along with changes in the BM and thymic stromal microenvironments, contribute to the early onset of thymus involution by 3MO of age.

It has previously been established that hematopoietic progenitors in the BM become myeloid-biased with age. For example, transplantation experiments revealed that HSCs from aged mice are biased towards a myeloid fate (Beerman et al., 2010; Busch et al., 2015; Min et al., 2004; Rossi et al., 2005; Young et al., 2016). Moreover, fate mapping studies have shown that MPPs from old mice transit more efficiently into common myeloid progenitors (CMPs) than CLPs (Busch et al., 2015). Nevertheless, the differentiation potential of lymphoid-biased HSCs on a per cell basis does not decline with age (Dorshkind et al., 2020; Montecino-Rodriguez et al., 2019). These prior studies assessed the decline in BM lymphoid potential between young (1-4MO) and much older mice (10-28MO). Since we find that ETP numbers decline by 3MO of age, we explored the possibility that a reduction in BM lymphoid progenitors contributes to the early decline in ETPs. Importantly, we found a striking decrease in the number of circulating TSPs, and the number and percentage of the BM progenitors Ly6d^-^ CLPs and Flk2^+^ MPPs at 3MO. Thus, diminished production and/or egress of T-lineage progenitors from the BM likely contribute to the early reduction in ETPs. Interestingly, a fetal-derived lymphoid-biased HSC subset persists in the BM until at least 2 weeks of age before being lost in adult mice (Beaudin et al., 2016). Also, embryonic-derived hematopoietic progenitors contribute to thymopoiesis through 7 weeks of age (Montecino-Rodriguez et al., 2018). These studies raise the possibility that the early decline in BM lymphoid progenitors and ETPs could reflect a reduced contribution of embryonically-derived hematopoietic progenitors. Of note, competitive heterochronic progenitor transfer assays, in which equivalent numbers of congenic 1MO and 3MO Flk2^+^CD27^+^ BM progenitors were injected into non-irradiated recipients of either age, showed no cell-intrinsic defect in the ability of 3MO lymphoid progenitors to seed the thymus or differentiate therein. To our knowledge, the present study is the first to show a substantial reduction in BM lymphoid progenitors and circulating TSPs by 3MO of age.

As lymphoid progenitor activity is regulated by niche factors in the thymus and BM microenvironments, age-associated changes in stromal cells could also play a role in the early decline of ETPs and BM lymphoid progenitors. In this regard, heterochronic BM transplantation experiments showed that successful T-cell reconstitution depends on the age of the recipient thymus rather than on the age of donor hematopoietic progenitors (Mackall et al., 1998; Zhu et al., 2007), consistent with the notion that thymic stromal support of thymopoiesis wanes with age. Alterations in the composition, organization and/or function of the thymus microenvironment could impair seeding and/or differentiation of TSPs, which enter the thymus on a periodic basis to occupy a limited number of stromal cell niches (Donskoy & Goldschneider, 1992; Foss et al., 2001; Zietara et al., 2015). Therefore, we asked whether the availability of functional TSP niches diminishes with age. Using multicongenic barcoding progenitor transfers, combined with mathematical modeling, thymuses from 1MO and 3MO old mice were found to contain a comparable number of available TSP niches. Thus, the early decline in ETP numbers is not a function of reduced TSP niche availability. To the contrary, our results showed an upward trend in the number of TSP niches with age. Previous investigations have reported evidence for a feedback loop in which key TSP niche factors and T-cell progenitor entry are restricted by the number of intrathymic T-cell precursors (Prockop & Petrie, 2004; Rossi et al., 2005; Zietara et al., 2015). Since we find a substantial reduction in the number of circulating TSPs and intrathymic ETPs as early as 3MO of age, it is possible that the observed increase in available TSP niches with advancing age is due, at least in part, to a decline in feedback from TSPs and/or ETPs.

After TSPs transition to the ETP stage, cTECs play a critical role in supporting their survival, proliferation and T-lineage commitment by providing niche factors, like DLL4, KITL and IL-7 (Han & Zuniga-Pflucker, 2021; Krueger et al., 2017; Lancaster et al., 2018). Given that the TEC compartment deteriorates during thymus involution, we considered the possibility that ETP niche factors become diminished with age. It is well established that Notch signaling plays a key role in T-lymphopoiesis (Chen et al., 2019; Han & Zuniga-Pflucker, 2021; Hozumi, Mailhos, et al., 2008; Koch et al., 2008; Tikhonova et al., 2019; Wilson et al., 2001). NOTCH1 is expressed on lymphoid progenitors in the BM and on ETPs in the thymus, while the critical Notch ligand DLL4 is expressed by vascular endothelial cells in both BM and thymus as well as on cTECs (Chen et al., 2019; Han & Zuniga-Pflucker, 2021; Tikhonova et al., 2019). Several observations in the present report suggest that diminished Notch signaling contributes to the early decline in BM lymphoid progenitors and ETPs between 1MO and 3MO of age. We found that lower levels of NOTCH1 are expressed on ETPs at 3MO of age, when *Dll4* expression is reduced on cTECs. Subsequent analysis of Notch signaling using the CBF-H2B-Venus Notch reporter strain (Nowotschin et al., 2013) demonstrated an age-related decline in Notch signaling in ETPs, as well as diminished Notch signaling in BM-resident CLPs. This decline may be due to deterioration of functional vascular and/or TEC niches. It is interesting to note that while the number of available TSP niches does not decline by 3 months, the ETP niche in the cTEC compartment may not be fully capable of supporting the progenitors that seed the thymus and enter the parenchyma, resulting in a lower number of detectable ETPs and downstream progeny. We did not find an increase in apoptosis of ETPs in the 3MO thymus, which might be expected in this scenario, but it is possible that clearance of ETPs undergoing apoptosis is quite rapid.

Altogether, our collective results indicate that an early age-associated decline in Notch signaling in T-cell progenitors could result in a reduced number of BM lymphoid progenitors, circulating TSPs and thymic ETPs at the outset of thymus involution. This potential link will be further evaluated in future studies. In summary, this study shows that an early age-associated decline in BM lymphoid potential and alterations in thymic stromal cells during the first 3MO of life synergize to initiate the earliest stages of thymus involution.

## Materials and Methods

### Mice

C57BL/6J (CD45.2, Thy1.2), B6.PL-Thy1a/CyJ 9 (CD45.2, Thy1.1), B6.SJL-*Ptprc*^*a*^ *Pepc*^*b*^/BoyJ (CD45.1, Thy1.2), C57BL/6-Tg(CAG-EGFP)1Osb/J (GFP) (Okabe et al., 1997) and Tg(Cp-HIST1H2BB/Venus)47Hadj/J (Notch-Venus reporter) (Nowotschin et al., 2013) mice were obtained from Jackson Laboratories and bred in-house. F1 mice for multicongenic barcoding experiments were bred in-house from CD45 congenic, Thy1 congenic, and EGFP strains (Fig. S2 B). C57BL/6J recipient mice for multicongenic barcoding experiments were obtained from the National Institute of Aging. All mice were maintained under specific pathogen-free conditions at the animal facilities at the University of Texas at Austin and the University of Texas MD Anderson Cancer Center, Smithville. All experimental procedures were performed in accordance with the Institutional Animal Care and Use Committees.

### Multicongenic barcoding progenitor transfer assay

BM cells were obtained from long bones of 4-6-week-old mice of 8 different congenic backgrounds (Fig. S2 B). Bones were crushed and single cell suspensions were prepared in FACS wash buffer (PBS+ 2% FBS), followed by RBC lysis (RBC lysis buffer, BioLegend). Cells were stained with lineage (Lin)-specific antibodies against CD11b, Gr-1, Ter-119, B220, CD19, CD3 and CD8 (all from BioXCell), and Lin^+^ cells were depleted with sheep anti-rat IgG immunomagnetic beads (Dynabeads, Invitrogen). Cells were then immunostained and Lin^-^ Flk2^+^CD27^+^ BM progenitors from F1 congenic strains were FACS purified (Fig. S2 A). Progenitors from each of the 8 congenic strains were mixed at equal ratios, and a total of 100,000 cells were retro-orbitally injected into non-irradiated 1MO, 3MO, 6MO and 12MO C57BL/6J recipient mice. Thymocytes were analyzed after 21 days by flow cytometry to distinguish the contributions of distinct congenic donor strains. 2.5 million events were recorded per recipient thymus, and successful thymic colonization was scored if a minimum of 40 cells of a given donor strain was detected. Quantification of niches was performed using a multinomial sampling algorithm (see Bayesian estimation of TSP niche numbers).

### Competitive heterochronic progenitor seeding assay

Lin^-^Flk2^+^CD27^+^ progenitors were isolated from the long bones of 1MO and 3MO congenic donor mice as described above. 1MO and 3MO progenitors were mixed at 1:1 ratio and 100,000 cells were retro-orbitally injected into 1MO and 3MO non-irradiated C57BL/6J recipient mice. After 21 days, donor chimerism was analyzed in the recipient thymi by flow cytometry.

### Flow cytometry

For analysis of thymocyte subsets, cell suspensions were prepared by manually dissociating thymuses in FACS wash buffer (FWB; PBS+ 2% bovine calf serum), and the cells were filtered through a 40μM cell strainer (Fisher). For isolating cTECs for qRT-PCR, thymi were isolated and cleaned of fat and connective tissue and placed in PBS. Thymic lobes were cut into small pieces and enzymatically digested as previously described (Seach et al., 2012). Briefly, 0.2mg of liberase TM (Sigma Aldrich) and 80 U of DNAse 1 (Sigma Aldrich) were used per thymus tissue for digestion. An enzymatic mixture containing Liberase TM and DNAse 1 in 2ml PBS were added to the thymic lobes and gently agitated in a 37°C shaking incubator for 10 minutes for a total of 4 digestion steps. At each digestion step, supernatant was collected into a 50ml tube containing FWB (PBS+ 2% bovine calf serum+ 5mM EDTA). Cells were centrifuged at 1250rpm for 5min at 4°C, and the pellet was resuspended in 5ml of FWB. Cells were depleted of thymocytes using anti-mouse CD45 Microbeads (Miltenyi Biotech) on the autoMACS® Pro Separator according to manufacturer’s recommendations. Enriched stromal cells were immunostained before FACS sorting cTECs. For quantification of circulating Lin^-^Flk2^+^CD27^+^ progenitors, following euthanasia, blood was collected immediately by making an incision in the right atrium and puncturing the left ventricle with a syringe. All possible blood from the mice was collected from the chest cavity by perfusing with 10mM of ethylenediaminetetraacetic acid/phosphate-buffered saline. RBCs were lysed with RBC lysis buffer (BioLegend) twice and white-blood cells were immunostained for flow cytometry. For quantification of BM lymphoid progenitors, long bones from each mouse were collected and crushed to obtain single-cell suspensions, prior to filtering through a 40μM cell strainer (Fisher) and lysing RBCs with RBC lysis buffer (BioLegend).

For preparation for flow cytometry, 6-10 million cells were immunostained with the fluorescently conjugated antibodies for 20 minutes on ice in FWB prior to washing in FWB and resuspending in FWB plus propidium iodide (Enzo) before flow-cytometric analysis. For intracellular staining of BrdU and cleaved caspase-3, cells were stained with antibodies for cell surface markers in FWB plus fixable viability dye (Zombie Red; BioLegend), then fixed and permeabilized using the BrdU APC kit (BD Biosciences) according to manufacturer’s protocol. Then the cells were stained with anti-BrdU and anti-cleaved caspase-3 fluorescently conjugated antibodies. Flow cytometry data was acquired on a BD LSRFortessa™ or and cells were FACS sorted on a BD FACSAria™ or BD FACSAria™ Fusion and analyzed using FlowJo software (V9.9.6 and V10.8.0, Tree Star Inc). A complete list of antibodies used for FACS analysis and sorting are provided in Table S1.

### BrdU Assay

Mice were weighed to determine the amount of BrdU to be injected per gram of body weight. 0.1mg of BrdU per gram of body weight was injected intra-peritoneally into 1MO and 3MO C57BL/6J mice. After 8 hours, thymocytes were isolated to assess BrdU incorporation by flow cytometry.

### cDNA preparation and quantitative PCR

FACS-purified cTECs from 1MO and 3MO C57BL/6J mice were resuspended in TRIzol reagent (Invitrogen) to isolate RNA. cDNA was prepared using the qScript cDNA synthesis kit (Quantobio) according to manufacturer’s instructions. qRT-PCR experiments were performed on the Viia7 Real-time PCR system (Thermo Fisher Scientific) using the following SYBR green and Taqman probes: DLL4 (SYBR green), actin (SYBR Green) and mouse alpha tubulin (Taqman). The primer sequences for DLL4 are as follows: forward primer, 5’-AGGTGCCACTTCGGTTACAC-3’; reverse primer, 5’-GGGAGAGCAAATGGCTGATA-3’. Gene expression levels were normalized relative to β-actin.

### Bayesian estimation of TSP niche numbers

For quantification of thymic niches, a Bayesian technique was implemented to estimate the number of niches present in recipient mice from the missing number of donors detected. Generalized linear models were used to test if the distributions of the estimated number of niches present in each age category declines with increasing age. General linear modeling was performed with Poisson response and a log link function with age as the only categorical covariate. Test statistic (χ^2^) was used to test the trend of number of niches over age groups. For estimation, the donor probabilities from the proportion of donor cells injected in each mouse was calculated. A posterior likelihood was computed using sampling from a multinomial distribution where L is the number of iterations, N is the number of niches and k is the number of distinct donors. This likelihood was then normalized to give a probability distribution for each likely value of missing donors. The maximum estimate of N that are drawn from the posterior distributions (MAP) were used as the most likely estimate of available niches. Batch to batch variation (i.e. replicate experiments) was used to test for potential bias due to batch effects. Regression analysis was used to assess if there are differences in available niches by age.

### Statistical analyses

Bayesian estimation to quantify the number of TSP niches was performed using R (The R Foundation for Statistical Computing). Statistical tests were done using Prism (V9.2.0; GraphPad Software), and significance was determined using an unpaired Student *t*-test, 1-way analysis of variance (ANOVA) with Kruskal-Wallis multiple comparisons test or 2-way ANOVA with Sidak’s multiple comparisons test, as indicated in the figure legends.

## Supplemental material

Fig. S1 shows the gating strategy for identification of thymocyte subsets and correlation of ETP cellularity with downstream thymocyte subsets. Fig. S2 shows gating strategies to sort Flk2+ CD27+ lymphoid progenitors from eight different congenic strains and to identify donor strain-derived thymocytes in recipient thymuses following transfer of the multicongenic progenitors. Fig. S3 shows that the Notch-Venus signaling reporter demonstrates faithful expression by immature thymocyte subsets. Table S1 lists all the antibodies used for flow cytometry analysis and FACS sorting.

## Data Sharing Statement

For original data, please contact

## Acknowledgements

We thank Janko Nikolich-Zugich, Nancy Manley, Marcel van den Brink, Jarrod Dudakov, Gregory Sempowski, Laura Hale, and Jen Uhrlaub for constructive feedback on experiments and manuscript preparation. This research was supported by a grant from the National Institutes of Health, P01AG052359, to L.I.R.E. and E.R.R.

## Authorship contributions

J.S., A.V., E.R. and L. E. designed the experiments and wrote the manuscript; J.S., A.V., H.S., E.P., performed experiments and analyzed data; B.L. and S.S. performed statistical analyses; A.K., E.R. and L.E. edited the manuscript.

