## Supplementary Material for "An early decline in ETPs reflects fewer pre-thymic progenitors and altered signals from the thymus microenvironment"

**Supplementary Figures**

Srinivasan et al.

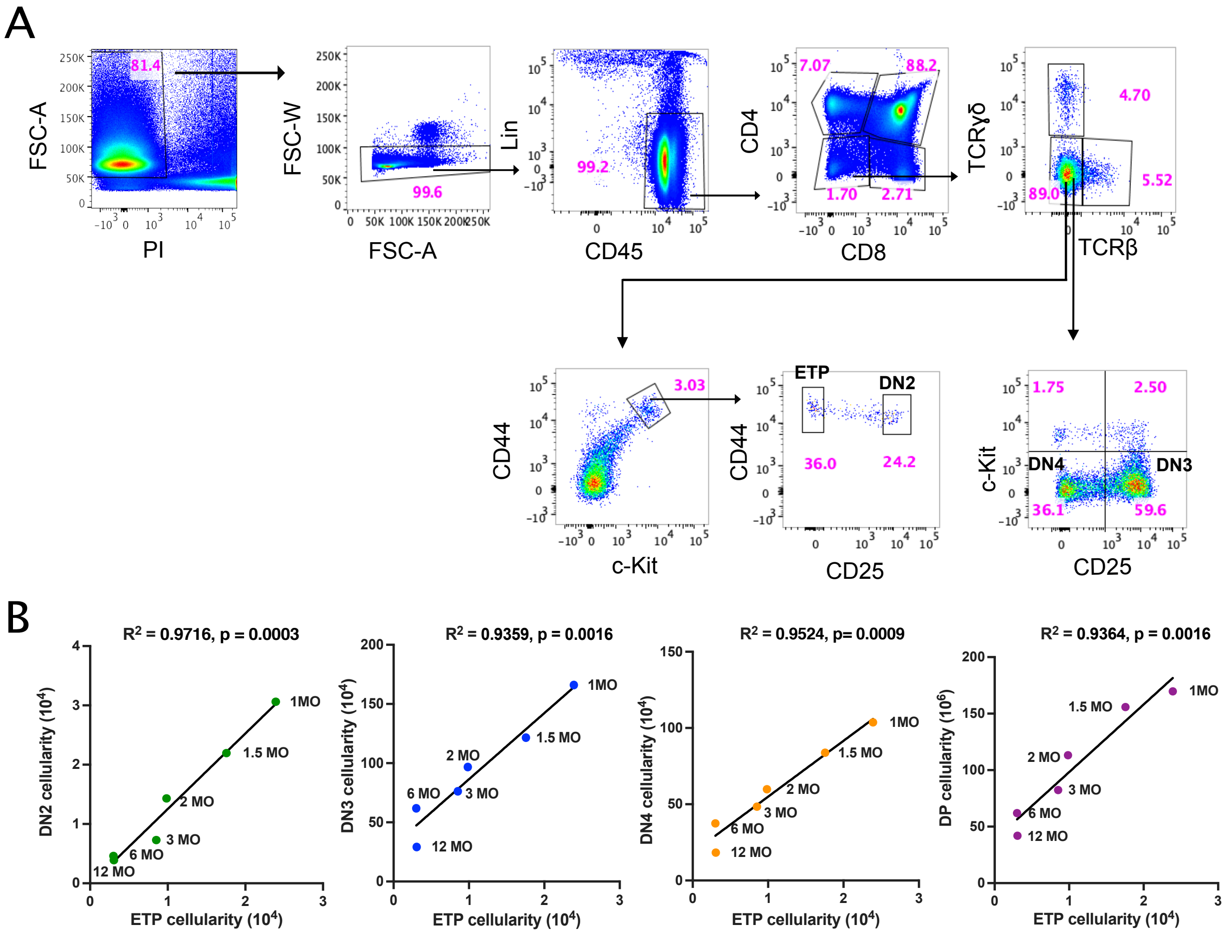


**Figure S1. Gating strategy for identification of thymocyte subsets and correlation of ETP cellularity with downstream thymocyte subsets.** (A) Representative flow cytometry plots showing gating strategy for ETPs (Lin$^-$ c-kit$^+$ CD44$^+$ CD25$^-$), DN2s (Lin$^-$ c-kit$^+$ CD44$^+$ CD25$^+$), DN3s (Lin$^-$ c-kit$^-$ CD25$^+$), DN4s (Lin$^-$ c-kit$^-$ CD25$^-$), as well as DN, CD4$^+$CD8$^+$ DP, CD4SP, and CD8SP thymocyte subsets. DN subsets were identified within Lin$^-$ CD4$^-$ CD8$^-$ TCR$\alpha\beta^-$ TCR$\gamma\delta^-$ thymocyte fraction. (B) Linear regression analysis of ETP cellularity and (from left to right) DN2, DN3, DN4 and DP cellularity at each age. $R^2$ = coefficient of correlation. Symbols represent data from an average of 7-14 mice at each age.

A

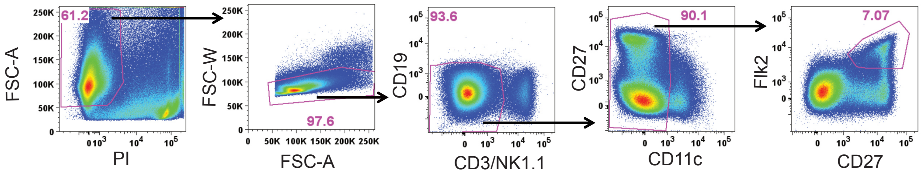


B

**Parental, recipient and eight F1 donor strains for the multicongenic barcoding experiment**

| Parental Strains | CD45.1; Thy1.2 | CD45.2; Thy1.2 | GFP+; CD45.2; Thy1.2 | CD45.2; Thy1.1 |
|---|---|---|---|---|
| CD45.1; Thy1.2 | (1) CD45.1; Thy1.2 | (2) CD45.1; CD45.2; Thy1.2 | (3) GFP+; CD45.1; CD45.2; Thy1.2 | (4) CD45.1; CD45.2; Thy1.1; Thy1.2 |
| CD45.2; Thy1.2 | | CD45.2; Thy1.2 (Recipient strain) | (5) GFP+; CD45.2; Thy1.2 | (6) CD45.2; Thy1.1; Thy1.2 |
| GFP+; CD45.2; Thy1.2 | | | | (7) GFP+; CD45.2; Thy1.1; Thy1.2 |
| CD45.2; Thy1.1 | | | | (8) CD45.2; Thy1.1 |

C

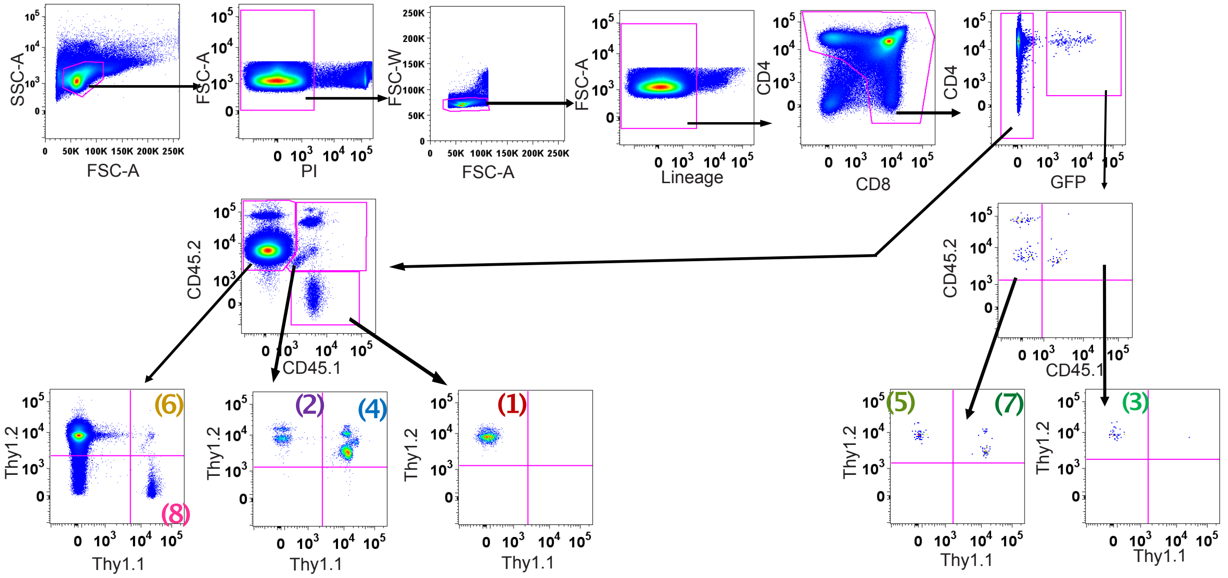


**Figure S2. Identification of donor strain-derived thymocytes in recipient thymuses following transfer of multicongenic progenitors.** (A) Representative gating strategy used for sorting multicongenic Flk2$^+$ CD27$^+$ progenitors. BM from all donors were depleted of mature lineages with antibodies against CD11b, CD11c, CD19, B220, Gr-1, NK1.1 and Ter-119. CD3$^+$, NK1.1$^+$ and CD11c$^+$ cells were excluded by gating and Flk2$^+$ CD27$^+$ progenitors were FACS sorted. (B) Eight distinct congenic F1 donor strains were generated by intercrossing CD45.1, CD45.2, GFP, and Thy1.1 strains. (C) Representative gating strategy to identify individual donor strains color-coded as in (B) 21d after i.v. transfer into non-irradiated 1MO, 3MO, 6MO and 12MO C57BL6/J recipient mice. The experiment was performed 4 times, with n = 10-12 recipients per age.

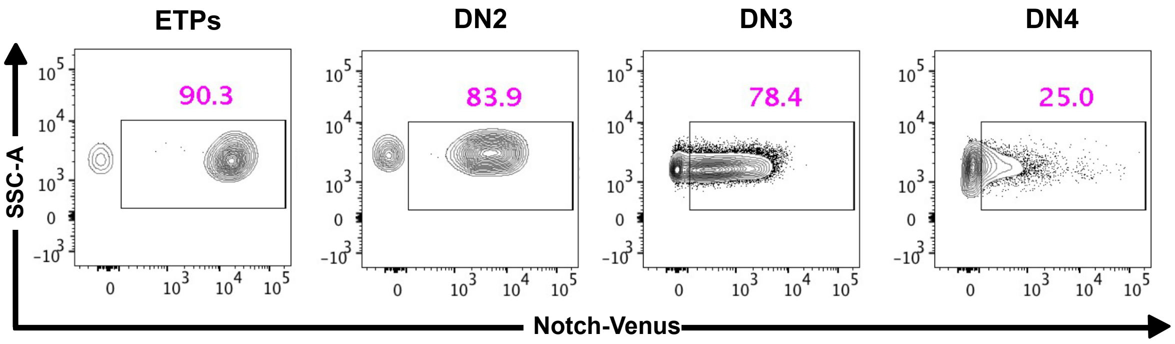


**Figure S3. Notch-Venus signaling reporter demonstrates faithful expression by immature thymocyte subsets**. Representative flow cytometry plots show the frequency of cells undergoing Notch signaling as well as Venus expression levels in the Notch-Venus reporter mouse strain for the indicated thymocyte DN subsets.

**Table S1: List of Antibodies**

| Target | Fluorophore | Manufacturer | Clone | Dilution |
|---|---|---|---|---|
| B220 | PE/Cy5 | eBioscience | RA3-6B2 | 1:50 |
| anti-BrdU | APC | BD Biosciences | | 1:100 |
| Cleaved Caspase-3 | BV421 | Cell Signaling Technologies | D3E9 | 1:100 |
| CD3 | PE/Cy5 | Biolegend | 145-2C11 | 1:50 |
| CD3 | PE/Cy7 | Biolegend | 145-2C11 | 1:200 |
| CD4 | BV510 | Biolegend | RM4-5 | 1:100 |
| CD4 | PE | BD Pharmingen | RM4-5 | 1:200 |
| CD4 | PerCP/Cy5.5 | Tonbo Biosciences | RM4-5 | 1:40 |
| CD8 | PE/Cy7 | eBioscience | 53-6.7 | 1:80 |
| CD8 | APC/Cy7 | Biolegend | 53-6.7 | 1:200 |
| CD11b | PE/Cy5 | Tonbo Biosciences | M1/70 | 1:50 |
| CD11c | PE/Cy5 | Biolegend | N418 | 1:50 |
| CD11c | PerCP/Cy5.5 | Tonbo Biosciences | N418 | 1:200 |
| CD25 | FITC | BD Pharmingen | 7D4 | 1:40 |
| CD25 | AF700 | Biolegend | PC61 | 1:800 |
| CD25 | Pacific Blue | Biolegend | PC61 | 1:50 |
| CD27 | APC | Biolegend | LG.3A10 | 1:200 |
| CD27 | PE/Cy7 | Biolegend | LG.3A10 | 1:200 |
| CD44 | APC | Biolegend | IM7 | 1:50 |
| CD44 | FITC | Tonbo Biosciences | IM7 | 1:400 |
| CD45 | PerCP/Cy5.5 | Invitrogen | 30-F11 | 1:100 |
| CD45 | BV510 | Biolegend | 30-F11 | 1:100 |
| CD45.1 | APC | Biolegend | A20 | 1:200 |
| CD45.2 | PE/Cy7 | Biolegend | 104 | 1:150 |
| c-kit (CD117) | APC/Cy7 | Biolegend | 2B8 | 1:40 |
| EpCAM | APC | Biolegend | G8.8 | 1:300 |
| Flk-2 (CD135) | PE | Biolegend | A2F10 | 1:200 |
| Flk2 (CD135) | APC | Biolegend | A2F10 | 1:50 |
| Gr-1 | PE/Cy5 | Biolegend | RB6-8C5 | 1:50 |
| IL-7R (CD127) | Biotin | Biolegend | A7R34 | 1:100 |
| Ly51 | PE | Biolegend | 6C3 | 1:300 |
| Ly6D | PE | Biolegend | 49-H4 | 1:200 |
| I-A/I-E (MHC-II) | APC/Cy7 | Biolegend | M5/114.15.2 | 1:200 |
| NK1.1 | PE/Cy5 | Biolegend | PK136 | 1:50 |
| NK1.1 | PE/Cy7 | Biolegend | PK136 | 1:200 |
| Notch1 | PE | Biolegend | HMN1-12 | 1:100 |
| Sca-1 | Pacific Blue | Biolegend | D7 | 1:200 |
| TCRβ | AF700 | Biolegend | H57-597 | 1:50 |
| TCRβ | PE | Biolegend | H57-597 | 1:40 |
| TCRγ/δ | PE | eBioscience | eBioGL3 | 1:40 |
| Ter-119 | PE/Cy5 | eBioscience | TER-119 | 1:50 |
| Thy1.1 | AF700 | Biolegend | OX-7 | 1:200 |
| Thy1.2 | BV510 | Biolegend | 30-H12 | 1:100 |
| UEA-1 | FITC | Vector Laboratories | | 1:500 |
| Fixable viability dye-zombie red | | Biolegend | | 1:1000 |
| | Qdot®605 | Invitrogen | streptavidin | 1:200 |
